# The Role of Secondary Chordae Tendineae in Mitral Valve Mechanics: A Benchtop Study

**DOI:** 10.64898/2026.07.20.736912

**Authors:** Roger Karl, Dominic P. Recco, Shannen Kizilski, Ibrahim Tharwat Abdelmoneim, Nicholas Kneier, Katrina Hon, C. Sören Bergt, Pedro J. del Nido, David Hoganson, Sandy Engelhardt, Peter Hammer

## Abstract

**Purpose:** Secondary chordae tendineae (CT) are frequently disregarded during mitral valve (MV) repair. However, their contribution to MV biomechanics remains poorly understood. We hypothesized that secondary CT play an important role in maintaining leaflet geometry during valve closure.

**Methods:** Eleven explanted porcine MVs were micro-CT scanned in a systolic configuration under three conditions: (1) physiological baseline with all secondary CT intact, (2) transection of the A2 strut CT, and (3) transection of all secondary CT. Billowing height (BH), coaptation height (CH) along the parasternal long-axis view, and coaptation area (CA) were quantified and compared between conditions. An increase in BH ***≥***2 mm from baseline was defined as pathological.

**Results:** Transection of the A2 strut CT increased BH by 3.1 mm, exceeding the pathological billowing threshold by 1.1 mm (***p_adj_* = 0.003**). CH decreased by 0.6 mm (***p_adj_* = 0.003**), whereas CA showed no significant change compared with baseline. Subsequent transection of all secondary CT further increased BH to 4.8 mm relative to baseline, corresponding to 2.8 mm above the billowing threshold (***p_adj_* = 0.003**). CH did not differ significantly, while CA decreased by 0.35 cm**^2^** compared with baseline (***p_adj_* = 0.029**). All values represent medians.

**Conclusion:** Secondary CT play a critical role in stabilizing MV leaflet geometry. Their transection resulted in increased leaflet billowing and reduced coaptation metrics associated with unfavorable MV repair outcomes. Preservation or reconstruction of secondary CT may therefore be an important consideration in MV repair strategies.

## 1 Introduction

Degenerative, or primary, mitral regurgitation (MR) arises from intrinsic or acquired structural abnormalities of the mitral valve (MV) apparatus, which comprises the leaflets, chordae tendineae (CT), papillary muscles, and annulus [46, 50]. The predominant etiology of primary MR is myxomatous degeneration, a non-inflammatory process characterized by extracellular matrix remodeling and structural weakening of the MV apparatus. Subsequent leaflet redundancy and/or CT rupture culminate in leaflet billowing and/or MV prolapse and consequent MR. Billowing is classically defined as systolic displacement of one or both mitral leaflets into the left atrium with or without MR, with leaflet excursion *<* 2 mm considered physiological and ≥ 2 mm beyond the annular plane diagnostic of pathologic billowing [7]. Chandra et al. identified billowing height as a strong predictor of underlying MV disease, demonstrating that a billowing height *>* 1 mm discriminated degenerative disease from normal valves. On the other hand, a coaptation line that is more than 6 mm below the annular plane in the ventricle is indicative of pathologic tethering, which is associated with poor MV function [5]. This delicate balance between normal curvature and pathological deformation highlights the biomechanical complexity of degenerative MV disease and underscores the challenge of achieving durable, optimal outcomes following mitral valve repair (MVr).

We hypothesize that suboptimal durability following MVr may, in part, reflect limitations of contemporary techniques—specifically, selective replacement of primary but not secondary CT [1, 37, 54]. Primary CT insert at the leaflet free edge, preventing tip prolapse (flail) beyond the annular plane and ensuring effective coaptation, whereas secondary CT attach to the ventricular surface of the leaflet body and are thought to primarily limit marginal billowing while preserving leaflet and ventricular geometry [34, 47, 57]. Of note, transection of secondary CT has been shown to improve leaflet mobility, reduce regurgitation, and avoid adverse ventricular remodeling for patients with functional (secondary) MR [40–43, 45]. However, the pathophysiological context of functional MR differs fundamentally from that of degenerative disease. In functional MR, left ventricular dilation induces leaflet tethering via the secondary chordae, thereby restricting leaflet motion. In degenerative MR, by contrast, secondary CT are typically not the primary structural abnormality, yet they are often still transected or left unaddressed. Furthermore, in congenital heart disease, billowing is a common feature resulting from incomplete formation or elongation of secondary chordae. This is can be seen in pediatric mitral and tricuspid valves but is most frequently observed in complex atrioventricular canal valves.

We contend that secondary CT play a more integral biomechanical role than previously appreciated, particularly in regulating leaflet billow and coaptation geometry—parameters that directly determine MR severity [13, 18, 20, 36, 49, 60, 64]. As an example, for a given annular and leaflet size, billowing results in more leaflet length being pulled out of the coaptation zone, effectively shortening coaptation height. Although this is clear from a biomechanics perspective, quantification of these relationships has not been sufficiently demonstrated. Notably, we were unable to identify any tissue-based studies within the past two decades that evaluate preservation or restoration of secondary CT as a strategy to enhance repair quality in degenerative MR. Accordingly, the primary objective of this study was to examine the association between secondary CT integrity and MV geometric parameters—specifically leaflet billow and coaptation—in an ex-vivo porcine model. By clarifying the mechanistic contribution of secondary chordae to valve geometry, this work aims to inform MV reconstruction strategies and determine whether secondary CT preservation or replacement may reduce recurrent regurgitation and the need for reoperation following MVr for primary MR.

## 2 Materials and Methods

### 2.1 Study Endpoints

The overarching goal of this work is to improve our knowledge about the role of secondary CT in the biomechanics of the MV. The primary endpoint of this study is billowing height. Secondary endpoints are coaptation height (CH), and coaptation area (CA).

#### 2.1.1 Billowing Height

Billowing height is conventionally defined as leaflet tissue bulging ≥ 2 mm above the annular plane. This definition is tailored to specific echocardiographic cross-sectional views, such as the parasternal long-axis (PLAX) view, and is shown in the appendix Figure 10.[7] In contrast, our micro-CT bench top setup enabled three-dimensional analysis of the MV with paired comparisons between baseline and altered valve states. Accordingly, for the purposes of this study, pathologic billowing was redefined as leaflet tissue bulging ≥ 2 mm above the baseline configuration. Therefore, the null hypothesis (*H*_0_) was that cutting the secondary CT does not result in a leaflet bulging exceeding 2 mm relative to baseline. Alternative hypothesis (*H*_1_): Cutting the secondary CT resulted in a leaflet bulging of more than 2 mm between the baseline and altered MV states and consequently in pathologic billowing.

#### 2.1.2 Coaptation Height

CH is defined as the length (one-dimensional) of contact between the anterior and posterior mitral leaflets perpendicular to the annulus plane and is shown in the appendix Figure 10. The study tested the effect of cutting secondary CT on CH. We hypothesized that cutting secondary CT increases billowing height and consequently decreases CH.

#### 2.1.3 Coaptation Area

In contrast to CH, CA is the area (two-dimensional) of contact between the anterior and posterior mitral leaflets. The study tested the effect of cutting secondary CT on CA. Similarly to CH, the coaptation area was expected to decrease as billowing increased. CA measurement provides a more holistic picture of the impact of CH changes along the entire coaptation zone.

### 2.2 Benchtop Model

To investigate the primary endpoint, eleven porcine MVs, comprising the mitral annulus, leaflets, CT, and papillary muscles, were harvested from adult-sized hearts obtained from LAMPIRE Biological Laboratories, Inc. (Pipersville, PA, USA). The MVs were sized prior to harvesting using an annuloplasty ring sizer set (Edwards Lifesciences Corporation, Irvine, CA, USA) to determine their anatomical dimensions.

The MVs were then sutured into custom-designed and 3D-printed frames, created using PTC Creo 11.0 software (Parametric Technology GmbH, Munich, BY, Germany) and fabricated from Clear Resin V4.1 on a Form 3 printer (Formlabs, Somerville, MA, USA). The frame design was tailored to fit within a micro-CT Scanner (Albira Si, Bruker Corporation, Billerica, MA, USA). The annular shape of the frame, including measurements of the annular circumference, anterior-posterior diameter, anterolateral-posteromedial diameter, commissural diameter, annular height, and annular angle, was based on the human normal control cohort data from a previous study [27], ensuring an anatomically accurate representation of the MV anatomy. On the atrial side of the frame, a rubber gasket (Dragon Skin FX-Pro, Smooth-On Inc., Macungie, PA, USA) sealed the interface between the frame and a vacuum chamber. The vacuum chamber, also fabricated from Clear Resin V4.1 on a Form 3 printer, served as a simulated left atrium. Additionally, two winches were positioned behind the papillary muscles, allowing for precise adjustment of their positioning and tension. The entire benchtop setup is shown in Figure 1.

**Fig. 1.**
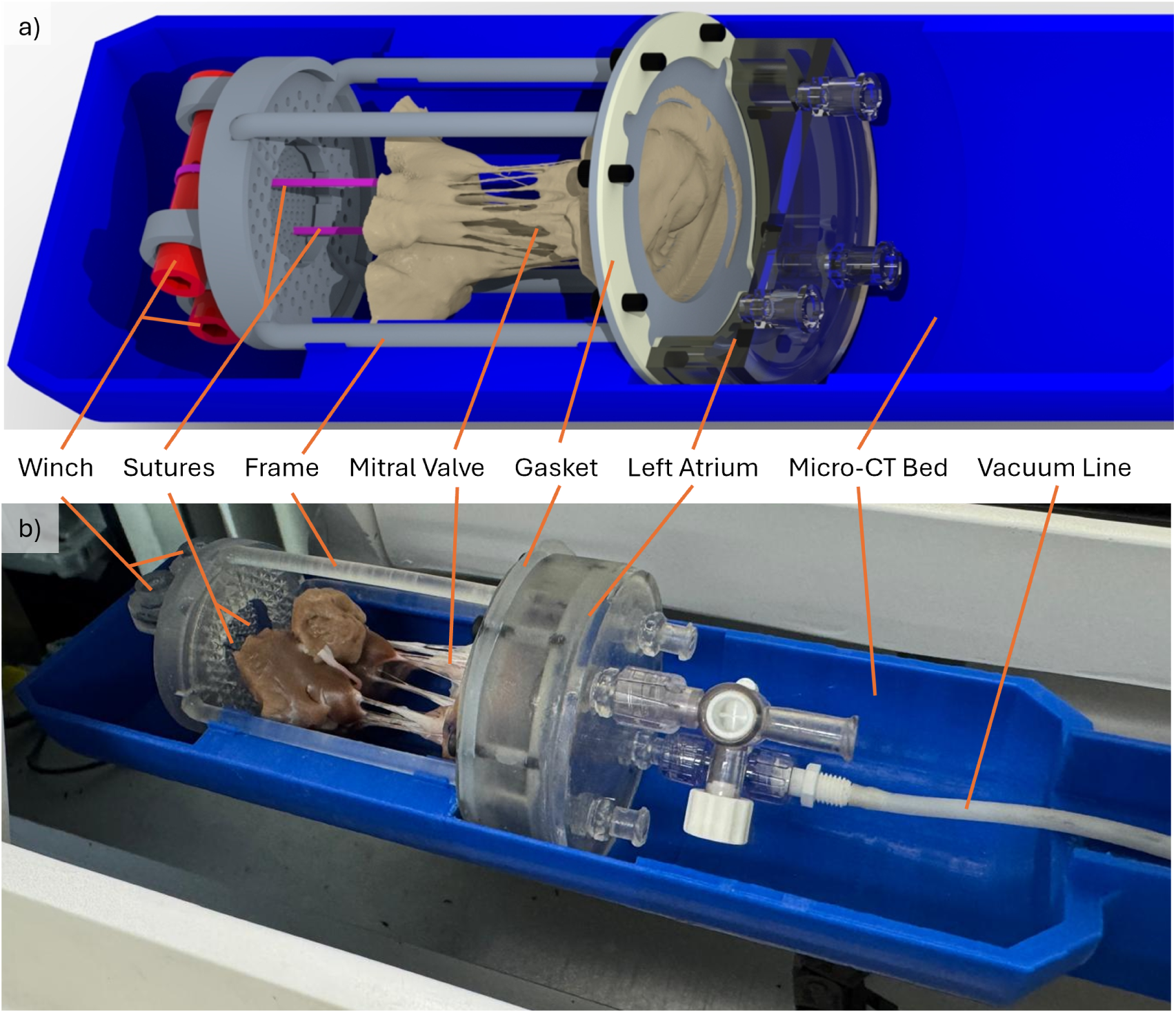
a) Computer aided design model of the benchtop setup. b) Photo of the benchtop setup with a sutured-in porcine valve

### 2.3 Study Design

Each sutured-in MV was placed inside a micro-CT scanner. To simulate average systolic conditions, a vacuum of 70 mmHg was applied to the left atrium, while the ventricular side of the valve was exposed to atmospheric pressure, resulting in a trans-mitral pressure differential of 70 mmHg. This pressure was chosen based on the physiological range of ventricular pressures during systole, which typically varies between slightly higher than left atrial pressure at the start of isovolumetric contraction (5-10mmHg) and a maximum pressure at peak systole (120 mmHg). Herein, the most commonly used MV annulus frame was 28 mm in inter-commissural diameter, which corresponds to a 17-year-old patient with a body surface area of 1.77 m², derived from our institution’s normalized data [14]. Based on this representative patient, we used a mean ventricular pressure of 77.5 mmHg and mean atrial pressure of 8 mmHg as reference points, resulting in an approximate trans-valvular pressure of 70 mmHg. To investigate the influence of secondary CT on biomechanics, each valve was evaluated under three conditions: (1) a physiological baseline condition with all CT intact, (2) transection of the strut CT at the A2 segment, and (3) transection of all secondary CT from the anterior and posterior leaflets.

To enhance visualization of the leaflet edge in the coaptation zone during micro-CT scanning, six micro titanium ligating clips (LIGACLIP™ Extra LT100, Ethicon, Inc., Raritan, NJ, USA) were applied, one at each of the leaflet segments, A1, A2, A3, P1, P2, and P3.

The experimental protocol consisted of four steps:

1. The papillary muscles were positioned to ensure anatomical closure and adequate leaflet coaptation while under vacuum.
2. A high-resolution micro-CT scan was performed using the following settings: high voltage (45 kV), high dose (400 µA), and a spatial resolution of 600 projections. The resulting voxel size was 0.125 mm x 0.125 mm x 0.125 mm.
3. The two strut CT, representing the thickest CT of the anterior leaflet, were transected, followed by a second micro-CT scan (Figure 2a & b).
4. Subsequently, all secondary CT (anterior and posterior leaflets) were transected, and a third micro-CT scan was acquired.

**Fig. 2.**
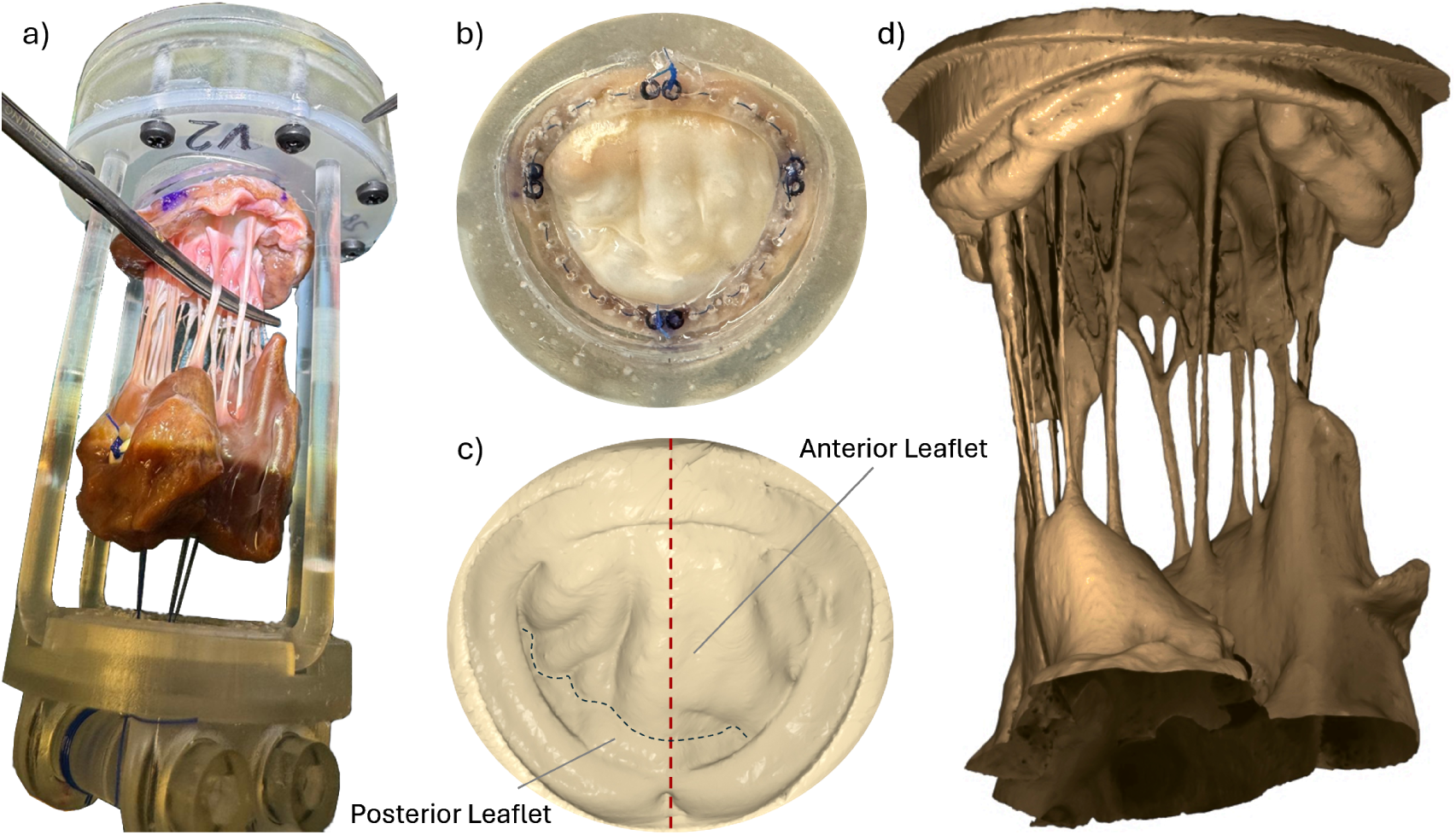
a) Subvalvular apparatus of the mitral valve (MV) displayed, with A2 strut chordae tenideae (CT) exposed. b) Atrial view of the MV under 70 mmHg vacuum. c) Atrial view of the micro-CT scanned, segmented, and smoothed MV, with the red dashed line indicating the parasternal long-axis view and the black dashed line following the line of coaptation between anterior and posterior leaflets. d) Subvalvular apparatus of the segmented MV

### 2.4 Measurements & Statistics

The micro-CT data were stored in DICOM format and imported into MITK (Medical Imaging Interaction Toolkit, German Cancer Research Center, Heidelberg, Germany). MVs were segmented in three steps. First, the majority of the surrounding frame was manually excluded using axial interpolation to remove structures not belonging to the model. Second, a threshold filter was applied to identify and visualize the ligating clips, using Hounsfield unit (HU) values from 1000 to the maximum of the dataset. Third, the MV tissue and its immediate surrounding structures were segmented using a threshold filter with HU values ranging from -600 to 600. This range represented a compromise between incomplete segmentation and excessive image noise. In some valves, larger segmentation gaps remained along the thin surface of the leaflet and were manually closed, as shown in Figure 2c & d. The MV tissue and the ligation clips were exported as two separate STL-files.

#### 2.4.1 Billowing Height

To investigate the primary endpoint, billowing height, the STL files were first smoothed using the Smooth modifier and manual sculpting tools in Blender 5.0 (Blender Foundation, Amsterdam, Netherlands). The processed STL files were then imported into CloudCompare 2.13.2. Rigid registration between baseline and altered MV models was performed using the Iterative Closest Point (ICP) algorithm. Following registration, the distance computation tool was applied to calculate the pointwise distance of each vertex in the baseline mesh to its nearest perpendicular point on the altered MV, as shown in Figure 3a - c. Maximum distance values were reported. Three one-tailed matched pairs Wilcoxon signed-rank tests were performed to compare the increase in billowing into the left atrium: (1) baseline versus the maximum distance after transected A2 strut CT, (2) baseline versus the maximum distance after all secondary CT were transected, and (3) transected A2 strut CT condition versus the all secondary CT transected MV condition. A Bonferroni correction was applied to adjust for multiple comparisons.

**Fig. 3.**
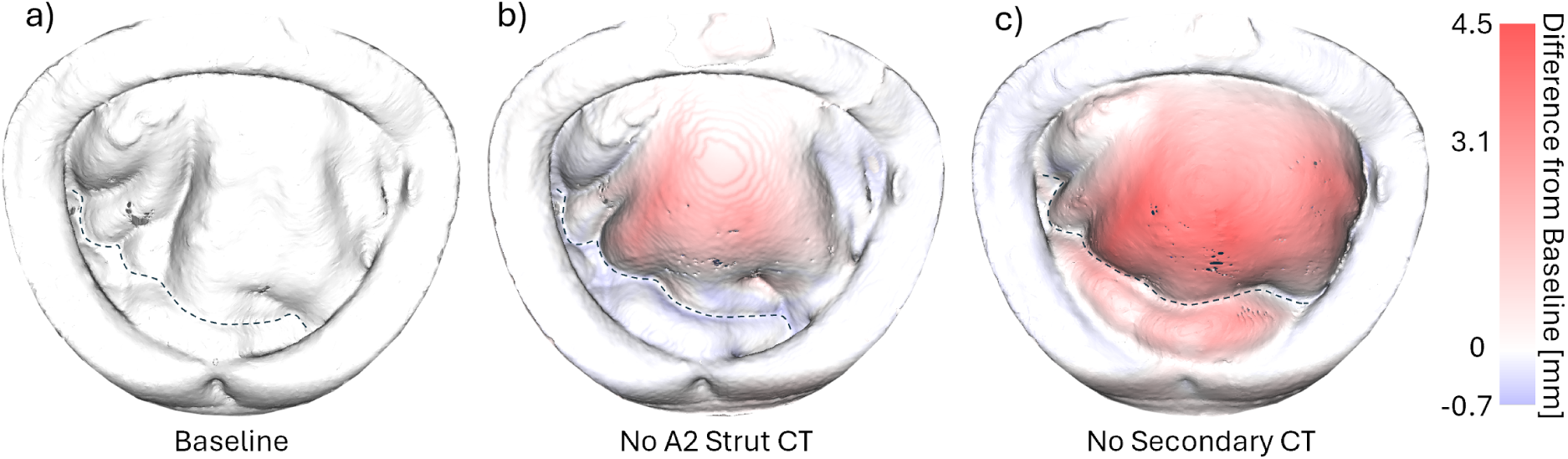
Representative heatmaps of billowing into the left atrium compared to a) baseline. Panel b) shows the condition with the two strut chordae tendineae (CT) of the A2 segment transected, resulting in a maximum billow of 3.1 mm. Panel c) illustrates transection of all secondary CT, with a maximum displacement of 4.5 mm. Positive values reflect displacement toward the atrium, while negative values reflect displacement toward the ventricle. Black dashed lines indicate the line of coaptation between the anterior and posterior leaflets

#### 2.4.2 Coaptation Height

CH was evaluated by measuring the shortest distance from the ligating clips in the A2/P2 segment to the highest coaptation point of the anterior and posterior leaflets in the PLAX view (Figure 4a). The STL files were imported into Creo Parametric to reconstruct the PLAX view and perform measurements. In the A2/P2 segment, two ligating clips were present, one on the anterior leaflet and one on the posterior leaflet. These clips define the ventricular (apical) end of the coaptation zone. As both leaflets are present only up to the level of the less apically positioned clip, measurements were taken from this clip to the atrial meeting point of the coaptation surface. CH was compared between the three different conditions of each MV. Figure 4b shows the PLAX view of these three conditions in one MV. Two two-tailed matched-pairs Wilcoxon signed-rank test were used. A Bonferroni correction was applied to adjust for multiple comparisons.

**Fig. 4.**
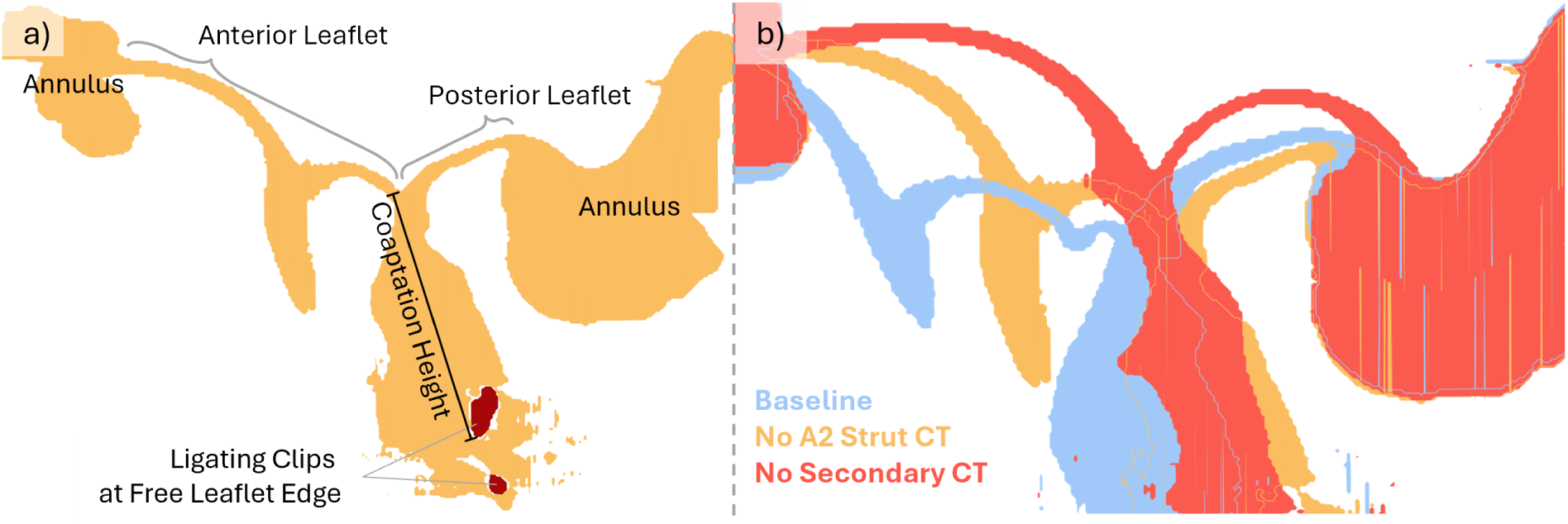
Parasternal long-axis view of the segmented mitral valve (MV). a) Illustration of coaptation height measurement from the most atrial point of leaflet coaptation (where the anterior and posterior leaflets meet) to the nearest ligation clip (red) at the leaflet free edge. b) Overlay of the MV under (blue) baseline condition, (orange) after transection of the two anterior strut chordae tendineae (CT), and (red) after transection of all secondary CT

#### 2.4.3 Coaptation Area

Direct measurement of the CA was not feasible due to its complex three-dimensional geometry within the MV apparatus. Therefore, changes in CA were estimated indirectly as the change in atrial-facing leaflet area from baseline to after cutting the secondary CT, based on the assumption that the total leaflet area (atrial leaflet area + two times coaptation leaflet area) remains constant (Figure 5). Consequently, the change in CA was assumed to be half of the change in the atrial leaflet area if both leaflets were affected equally. The atrial leaflet surfaces were segmented in Blender, and their surface areas were quantified using the 3D Print Toolbox add-on. Two two-tailed matched-pairs Wilcoxon signed-rank tests were performed between the test conditions and baseline. A Bonferroni correction was applied to adjust for multiple comparisons.

**Fig. 5.**
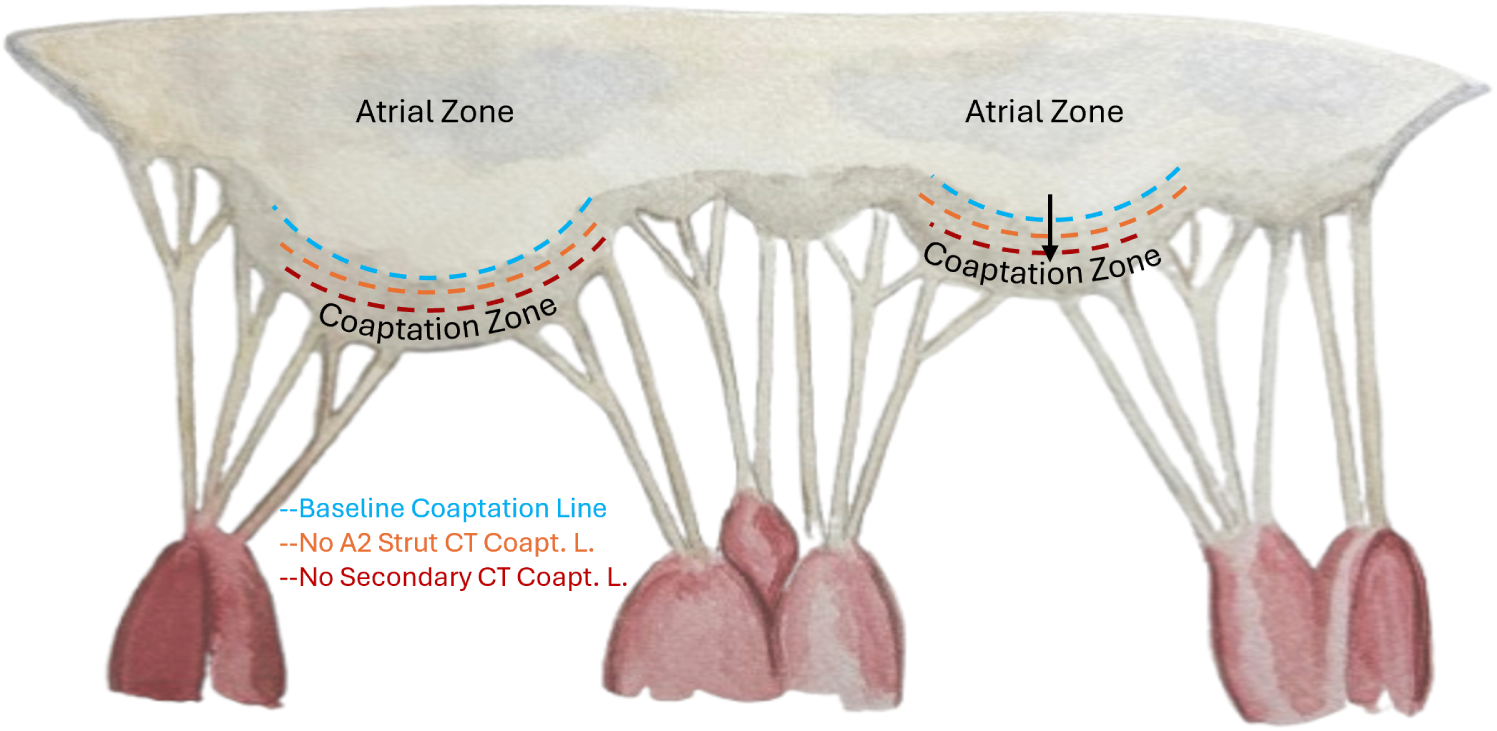
Unfolded view of the mitral valve highlighting the atrial zone and the coaptation zone. Dashed lines indicate the hypothesized shift of the coaptation line between the atrial and coaptation zones following transection of the chordae tendineae (CT). Redrawn and adapted from [11] Figure 5-15

#### 2.4.4 Statistical Analysis

Normality was assessed using the Shapiro–Wilk test. Four parameters were normally distributed: billowing height (baseline vs. no A2 strut CT; baseline vs. no secondary chordae tendineae), coaptation height (baseline vs. no secondary chordae tendineae), and coaptation area (baseline vs. no A2 strut chordae tendineae). Two parameters were not normally distributed due to a single outlier: coaptation height (baseline vs. no A2 strut chordae tendineae) and coaptation area (baseline vs. no secondary chordae tendineae). Statistical analyses and plots were performed using MATLAB R2025b (The MathWorks, Inc., Natick, MA, USA).

#### 2.4.5 A priori Power Study

After inclusion of the first five valves, a power analysis was performed using G*Power 3.1.9.7 based on the primary endpoint, CH. A one-tailed Wilcoxon signed-rank test for dependent samples (matched pairs) was applied. The effect size (Cohen’s d = 2.0) was calculated based on these initial five valves (mean difference = 1.2 mm; standard deviation of the differences = 0.6 mm). Assuming a normal parent distribution, a significance level of *α* = 0.01 and a desired power of 0.95, the analysis yielded a required total sample size of 8 valves.

### 2.5 Physiological Baseline

To ensure a physiological condition of the baseline valve, the maximal systolic displacement of leaflet tissue towards the atrium (billowing height) and the maximal systolic displacement of coaptation line towards ventricle (tenting height), were measured at the PLAX. A billowing height *<* 2 mm and a tenting height of *<* 6 mm compared to the annular plane are considered physiologically normal. Tenting height and billowing height are shown in the appendix Figure 10.

## 3 Results

Eleven excised porcine MVs underwent two-stage secondary CT cutting and micro-CT scanning to investigate the influence of secondary CT on the biomechanics of the MV.

### 3.1 Billowing Height

Transection of the A2 segment strut CT resulted in billowing into the left atrium ranging from 2.2 to 4.4 mm compared to baseline, with a median of 3.1 mm, corresponding to a median of 1.1 mm beyond the 2 mm pathological billowing threshold (Figure 6). Transection of all secondary CT further increased billowing by 0.5 to 2.5 mm with a median increase of 1.4 mm. In total, transecting all secondary CT led to billowing between 2.7 and 6.8 mm, with a median of 4.8 mm, corresponding to 2.8 mm beyond the 2 mm pathological billowing threshold. Consequently, as hypothesized, cutting secondary CT leads to tissue displacement of more than 2 mm for both transection of the A2 strut CT alone and transection of all secondary CT (*p_adj_*= 0.003 for each). Transection of all secondary CT increased billowing height significantly compared to transection of the A2 strut CT alone (*p_adj_*= 0.003).

**Fig. 6.**
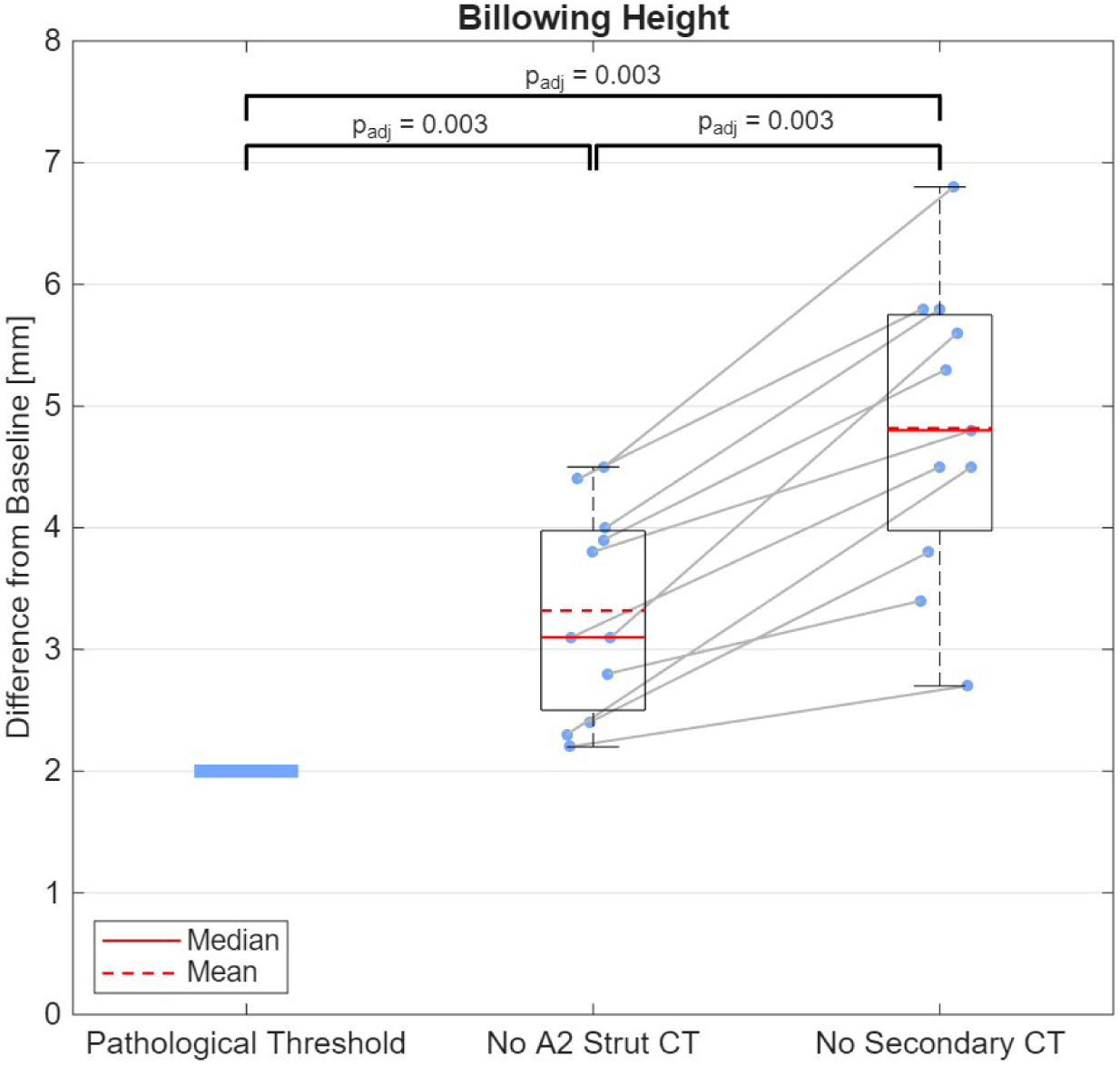
Maximum differences in billowing height from baseline were measured for each of the *n* = 11 valves with the strut chordae tendineae (CT) of the A2 segment transected and all secondary CT transected. Groups were compared to the mitral valve pathological billowing threshold, defined as a 2 mm offset from baseline. Statistical significance was assessed using one-tailed matched-pairs Wilcoxon signed-rank tests with Bonferroni correction for multiple comparisons

### 3.2 Coaptation Height

The median baseline CH for the valves in this experimental setup was 8.8 mm, ranging from 5.8 mm to 14.3 mm. Transection of the strut CT resulted in a reduction of CH along the PLAX view ranging from -0.1 to -5.2 mm, with a median decrease of -0.6 mm, indicating a significant decrease (*p_adj_*= 0.003), shown in Figure 7. Transection of all secondary CT resulted in a non-significant change ranging from -2.6 to +5.0 mm, with a median increase in CH of +0.9 mm (*p_adj_*= 0.126). Compared to the baseline condition, transection of all secondary CT led to a median increase in CH of 0.6 mm, with values ranging from - 5.1 mm (decrease) to +4.5 mm (increase) (*p_adj_* = 1). Since transecting all secondary CTs did not result in a significant change in CH, contrary to our expectations, we conducted a more detailed analysis. Across different locations, a reduction in CH was observed in 9 of the 11 MVs examined. Moreover, analysis of the correlation between individual leaflet billowing height and coaptation line movement revealed that transecting all secondary CTs causes the coaptation area in the A2/P2 segment to migrate towards the left atrium, thereby preserving CH. The corresponding analyses are presented in Appendix A.1.

**Fig. 7.**
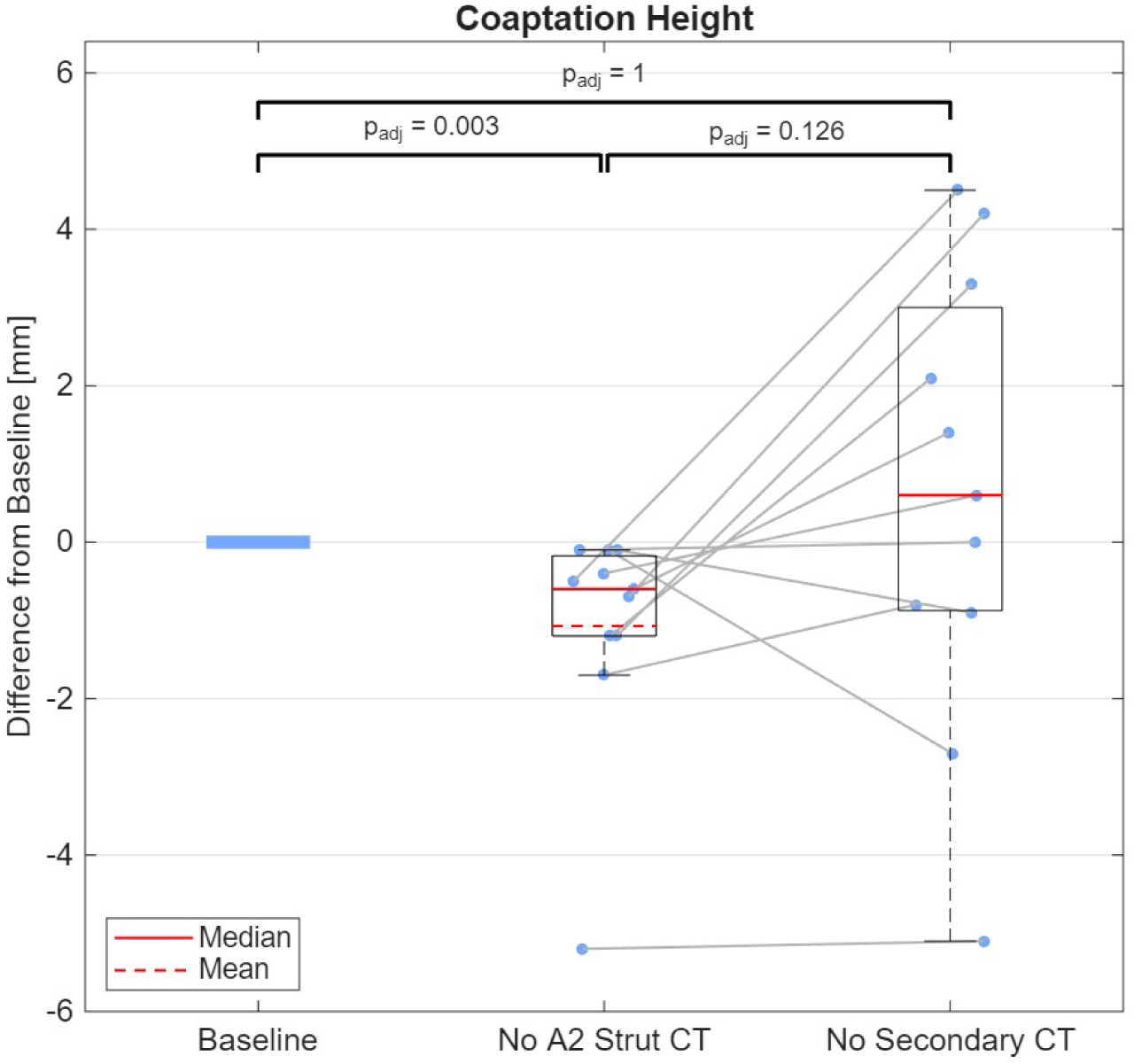
Coaptation height differences from baseline were measured at the parasternal long-axis view for each of the *n* = 11 valves with the strut chordae tendineae (CT) of the A2 segment transected and all secondary CT transected. Groups were compared to baseline. Statistical significance was assessed using two-tailed matched-pairs Wilcoxon signed-rank tests with Bonferroni correction for multiple comparisons

### 3.3 Coaptation Area

Transection of the strut CT resulted in a change in atrial leaflet area ranging from -0.13 to +0.68 cm^2^, with a median of +0.07 cm^2^. Consequently, the change in CA ranged from -0.38 to +0.06 cm^2^, with a median of -0.04 cm^2^ and is not significant (*p_adj_* = 0.352). Transection of all secondary CT led to changes, compared to transection of the A2 strut CT alone, in atrial leaflet area ranging from -0.21 to +3.50 cm^2^ with a median increase of 0.41 cm^2^, resulting in a change in CA ranging from -1.75 to +0.11 cm^2^ with a median decrease of -0.21 cm^2^, indicating a significant decrease (*p_adj_*= 0.029). Transection of all secondary CT compared to the baseline led to a change in atrial leaflet area ranging from -0.27 to +4.18 cm^2^, with a median of +0.69 cm^2^, corresponding to a significant (*p_adj_*= 0.029) change in CA ranging from -2.09 to +0.14 cm^2^ with a median of -0.35 cm^2^.

CA changes correspond inversely to change in atrial leaflet area, with values ranging from 50% when both leaflets are equally affected to 100% when only a single leaflet is involved. The values reported and shown in Figure 8 represent the inverse 50% of the atrial leaflet area, as a conservative estimation. Estimations of more than 50% would lead to a further decrease in CA.

**Fig. 8.**
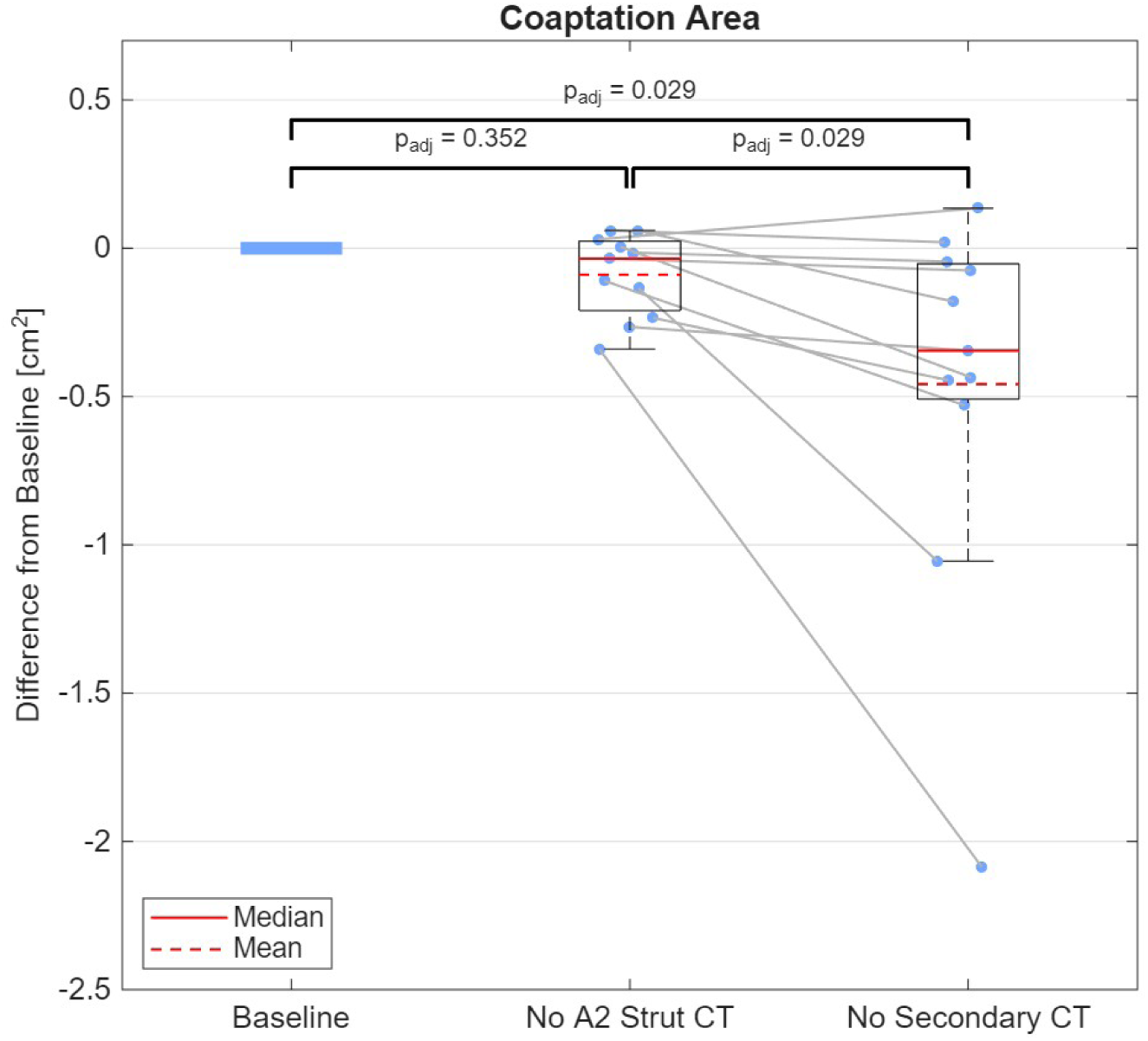
Coaptation area differences from baseline were calculated by comparing atrial leaflet areas for each of the *n* = 11 valves with (a) the strut chordae tendineae (CT) of the A2 segment transected and (b) all secondary CT transected. Groups were compared to baseline. Statistical significance was assessed using two-tailed matched-pairs Wilcoxon signed-rank tests with Bonferroni correction for multiple comparisons

### 3.4 Physiological Baseline

The tenting height in all valves at baseline was between -1.2 mm and +4.9 mm with a median of +1.2 mm. No MV had a tenting height considered abnormal (*>* 6 mm). Baseline billowing height ranged from +0.1 to +4.5 mm, with a median of +0.8 mm. Two MVs exhibited billowing greater than the +2 mm pathological billwoing threshold, considered abnormal, at baseline.

## 4 Discussion

### 4.1 Impact of Secondary Chordae Transection on Leaflet Geometry

Herein, we demonstrate the critical contribution of secondary CT to MV biomechanics through controlled ex-vivo experimentation. Porcine MVs were selected given their close anatomic similarity to human valves, as established by prior morphometric analyses [2]. An atrially positioned vacuum system was employed to optimize visualization of the sub-valvular apparatus, thereby enabling precise assessment of leaflet apposition and controlled transection of secondary CT. High-resolution micro-CT was used to facilitate detailed segmentation and sub-millimeter quantification of all measured endpoints. To enable direct within-specimen comparisons, paired experiments were performed in which each of the 11 valves underwent serial assessment under three conditions: (1) baseline, (2) following transection of strut CT at the A2 segment, and (3) following transection of all secondary CT. All outcomes were analyzed relative to baseline. Transection of both A2 strut CT and all secondary CT resulted in significant increases in billowing height, with median increases of 3.1 mm and 4.8 mm, respectively. Notably, both conditions exceeded the 2 mm threshold consistent with pathologic billowing, underscoring the biomechanical relevance of secondary CT integrity. Moreover, complete secondary CT transection produced significantly greater billowing than isolated strut chord transection, suggesting a cumulative structural role of secondary CT in maintaining leaflet restraint. With respect to coaptation-related metrics, isolated strut CT transection resulted in a modest but significant reduction in CH (median -0.6 mm). In contrast, transection of all secondary CT yielded CH values comparable to baseline and significantly greater than those observed after strut chord transection alone. When evaluating the coaptation line, we observed a clear relationship between the degree of posterior leaflet billowing and atrial displacement of the coaptation line. Following transection of all secondary CT, both leaflets shifted atrially, resulting in coaptation occurring at or above the annular plane. Because billowing involved both the anterior and posterior leaflets, CH remained comparable to baseline, albeit at a more atrialized position. Despite this preserved coaptation, such a configuration would likely confer a limited functional reserve, with minimal tolerance before the onset of clinically significant regurgitation. Because CH could only be measured at the A2/P2 location with high confidence due to model constraints, we hypothesized that this apparent preservation of CH reflects diffuse geometric redistribution of leaflet tissue rather than true maintenance of coaptation architecture. To further interrogate coaptation integrity, atrial leaflet area was evaluated as a surrogate for CA. While CA remained similar to baseline following strut CT transection, it decreased significantly by up to 2 cm^2^ after transection of all secondary CT, indicating compromised global coaptation despite preserved focal CH measurements. These findings were supported by reductions in CH in 9 out of 11 valves along the zone of coaptation but in different locations. Collectively, these findings highlight the integral role of secondary CT in preserving leaflet restraint and coaptation geometry. These findings also illustrate that adding primary edge chordae may not correct pathologic billowing, as in these leaflets, all primary CT were left intact.From a clinical perspective, these data support strategies aimed at preserving or reconstructing secondary CT during MVr for primary MR, as disruption of these structures may predispose a patient to recurrent regurgitation and subsequent reoperation.

### 4.2 Clinical Context: Current Mitral Valve Repair Strategies & Limitations

Two principal surgical strategies are employed for MVr in degenerative MR: chordal replacement and leaflet resection. Both approaches seek to reduce excessive leaflet billow and restore adequate coaptation length. Chordal replacement involves implantation of artificial CT (neo-chordae), most commonly fashioned from ePTFE, which are secured to the leaflet free edge and anchored to the native papillary muscle or ventricular myocardium to prevent persistent prolapse [44, 54]. In contrast, leaflet resection removes redundant leaflet segments to eliminate billowing tissue and re-establish leaflet alignment. Despite their procedural differences, these techniques have generally demonstrated comparable outcomes with respect to survival, left ventricular function, need for reintervention, and MR recurrence [10, 52, 61]. Some studies suggest potential advantages of neo-chordae implantation, citing improved preservation of leaflet mobility and coaptation [38, 51, 52]. Supporting this concept, Bennati et al. demonstrated in CFD simulations that neo-chordae repair produced more physiological transvalvular flow patterns compared with resection [6]. Nevertheless, durability after chordal replacement remains sub-optimal. In adult populations, recurrent MR has been reported in up to 28% of patients within 10 years following repair with neo-chordae [12, 15–17, 23, 25, 31, 33, 52, 56], with reoperation rates approaching 16% at 10 years. Although data in pediatric cohorts are comparatively limited [25, 32, 48], outcomes appear even less favorable, with recurrent MR reported in 17–30% and reintervention rates of 11–33% at 5–10 years post-repair [4, 8, 26, 28, 36, 63]. Similarly to adults, in pediatrics, primary chordae replacement and annuloplasty techniques are often utilized, with resection being less commonly employed. In combination, these two techniques shift more of the leaflet down into the coaptation zone. However, none of those techniques primarily addresses billowing. Our results unveil an opportunity to directly address leaflet billow and more efficiently and effectively increase CH/CA. Collectively, these data underscore a persistent gap between immediate procedural success and long-term durability.

### 4.3 Predictors of Recurrent Mitral Regurgitation & Mechanistic Framework

In response to these suboptimal long-term outcomes, prior investigations have sought to identify predictors of recurrent MR and MV reintervention following repair. Independent risk factors for structural valve deterioration include greater than mild residual MR at hospital discharge [23, 36, 56, 63], a lower number of implanted neo-chordae [21, 31], reduced coaptation length (*<*5–10 mm) [21, 60], and greater billow or prolapse extent [36, 49, 64]. However, these variables are unlikely to act in isolation. Coaptation length and billow and prolapse severity all strongly correlate with MR grade [13, 18, 20, 36, 49, 60, 64], suggesting a shared mechanistic pathway. From a biomechanical perspective, this interdependence is intuitive. Early systolic leaflet billowing promotes annular expansion, followed by progressive leaflet deformation and free-edge prolapse [3]. In the presence of MR, left ventricular dilation further displaces the papillary muscles, resulting in secondary leaflet tethering that exacerbates regurgitation [49]. This maladaptive positive feedback loop underscores the central surgical objective during MV repair: to reduce pathologic billow while restoring sufficient coaptation. Physiological billowing is typically characterized by leaflet excursion *<*2 mm beyond the annular plane, and normal coaptation height in healthy valves ranges from 5–10 mm [34, 35]. Notably, the coaptation heights associated with favorable postoperative outcomes are strikingly similar, generally reported between 4 and 11 mm [19–21, 39, 60, 64]. Despite this substantial body of literature, the optimal approach to achieve durable MVr is not well defined, inspiring the current study.

### 4.4 Integration with Computational Literature on Secondary Chordae

Although few groups have experimentally evaluated the biomechanical consequences of secondary CT transection ex-vivo, a substantial body of in-silico work has examined their structural and functional role. Computational studies by Prot, Toma, and Caballero have modeled chordal rupture and assessed its effects on leaflet stress distribution, papillary muscle loading, and valve geometry [9, 22, 53, 58]. Prot et al. reported that secondary CT bear approximately 31% of the load transmitted to the papillary muscles; removal of anterior strut chordae nearly doubled tension in the anterior primary CT and significantly increased anterior leaflet billowing [53]. Using fluid–structure interaction modeling, Caballero et al. demonstrated similar geometric and stress alterations [9]. Hammer et al. found that omission of secondary CT resulted in a peak atrial displacement of the MV leaflets by ∼7 mm [22]. Toma et al. further showed that rupture of secondary CT produced a greater regurgitant orifice area than rupture of primary CT [58]. Additional modeling approaches incorporating secondary CT include topology optimization and patient-specific finite element models [29, 59]. Reconstruction strategies for MVr have also been explored computationally[24, 55]. Collectively, these investigations consistently support a stabilizing role for secondary CT in load sharing, leaflet restraint, and stress redistribution. Our experimental findings complement and extend this in-silico literature by providing direct ex-vivo evidence that secondary CT integrity significantly influences billowing height, CH, and CA.

### 4.5 Clinical Implications for Mitral Valve Repair & Partial Heart Transplantation

One direct clinical implication of this work relates to both adult and pediatric MV reconstruction. Given the heterogeneity of MV disease and the breadth of surgical techniques required to address these complex pathologies, physical simulation platforms are often essential for both trainees education and patient-specific procedural planning[30]. The demonstrated influence of secondary CT can inform the design of ex-vivo and in-vitro dynamic simulators for surgical and transcatheter MV interventions, enabling more physiologically accurate training environments and improved personalization of repair strategies. At a more complex end of the reconstructive spectrum, these findings also have implications for atrioventricular valve partial heart transplantation, an approach that remains in the early stages of clinical adoption [62]. In current practice, surgeons generally replace only primary chordae attached to the leaflet free edge. In light of the demonstrated biomechanical contribution of secondary CT, our findings suggest that reconstruction strategies limited to primary chordal replacement incompletely restore native MV geometry. While it is neither practical nor necessary to recreate the full complement of secondary CT present in the normal human valve [2], our data indicate that the two strut CT alone account for approximately 65% of the observed increase in billowing height following complete secondary CT transection and contribute to measurable reductions in coaptation height. Selective incorporation of strut neo-chordae, therefore, may represent a pragmatic compromise—enhancing leaflet restraint and geometric fidelity without substantially prolonging operative time or cardiopulmonary bypass exposure. Such strategies may improve restoration of physiological leaflet geometry, enhance repair durability, and ultimately translate into superior long-term patient outcomes.

### 4.6 Limitations

The main limitations of this study relate to geometric aspects of the experimental setup. The shape and dimensions of the model may not fully replicate physiological conditions. Suturing the porcine valve into the fixture may have introduced minor misalignment, and the positioning of the papillary muscles as well as force transmission may differ from in vivo physiology. Although the system was designed to approximate physiological anatomy as closely as possible, these factors may have resulted in non-physiological geometry, potentially influencing leaflet coaptation or promoting billowing. However, owing to the matched-pair study design, such systematic effects are expected to be consistent across conditions and therefore unlikely to substantially affect the comparative results. Two of the baseline MVs exhibited abnormal geometry due to initial billowing and were retained in the analysis, as the progression of billowing and coaptation in these specimens was consistent with that observed in valves with normal baseline geometry. Their inclusion is therefore unlikely to have introduced a systematic bias into the comparative results. In addition, visualization of the coaptation zone was limited. This region is geometrically complex and characterized by anisotropic leaflet thickness and X-ray attenuation properties similar to adjacent soft tissues and chordae tendineae, resulting in low contrast in micro-CT imaging. Consequently, precise quantification of coaptation height or area remains challenging. To our knowledge, no more accurate or reliable method is currently available for assessing this region under the given experimental conditions.

### 4.7 Conclusions

Secondary chordae tendineae play an important role in stabilizing MV leaflet geometry. In this bench-top model, transection of secondary CT increased leaflet billowing and impaired coaptation metrics associated with MR. These findings suggest that secondary chordae contribute significantly to physiological valve mechanics. Preservation or reconstruction of secondary chordae may therefore represent an important consideration during MVr.

## Acknowledgements

The authors gratefully acknowledge Mara Thompson for valuable discussions and advice on the geometrical analysis of the 3D scanned data, as well as for guidance in the use of 3D modeling and analysis tools. Furthermore, the authors sincerely thank Dr. Jessica Körner for her contribution in creating the artwork for Figure 5.

## A Appendix

### A.1 Additional Analysis of Coaptation Height

Complete secondary CT transection did not significantly reduce coaptation height (CH) at the PLAX view (median change: +0.6 mm; only 4/11 MVs showed a reduction). We hypothesize this apparent preservation reflects diffuse leaflet reorientation rather than maintained coaptation architecture. To test this, CH was assessed at secondary ligating clip positions, revealing reductions in 5 additional MVs (9/11 total). However, due to irregular clip alignment outside the standardized PLAX view, these supplementary measurements were excluded from the main results.

Furthermore, we hypothesized that transecting all secondary CT would produce a pronounced shift of the MV apparatus, particularly at the coaptation zone in the A2/P2 region. To investigate this, we compared the billowing of the anterior leaflet, posterior leaflet, and the displacement of the coaptation line (= tenting height) at the A2/P2 level towards the left atrium, as shown in Figure 9. Selective transection of the A2 strut CT resulted in a median anterior leaflet billowing of 3.1 mm (range: 2.2–4.4 mm) toward the left atrium. The posterior leaflet exhibited a comparatively modest median shift of 0.3 mm toward the left atrium (range: 0.1 to 1.3 mm), which was anticipated given that only the anterior strut CT were transected. The coaptation line shifted a median of 0.5 mm toward the left atrium (range: -0.8 to 1.7 mm). Following transection of all secondary CT of both the anterior and posterior leaflets, the anterior leaflet demonstrated a median billowing of 4.6 mm (range: 2.4–6.8 mm), while the posterior leaflet billowed a median of 2.7 mm (range: 1.7 to 4.5 mm). The increase in displacement for both leaflets was expected, as a larger number of CT were transected at both leaflets. The coaptation line shifted a median of 2.0 mm toward the left atrium (range: -0.1 to 5.0 mm), closely mirroring the magnitude of posterior leaflet billowing, suggesting that either posterior leaflet or potentially rather bileaflet billowing is a primary determinant of coaptation line migration under these conditions.

**Fig. 9.**
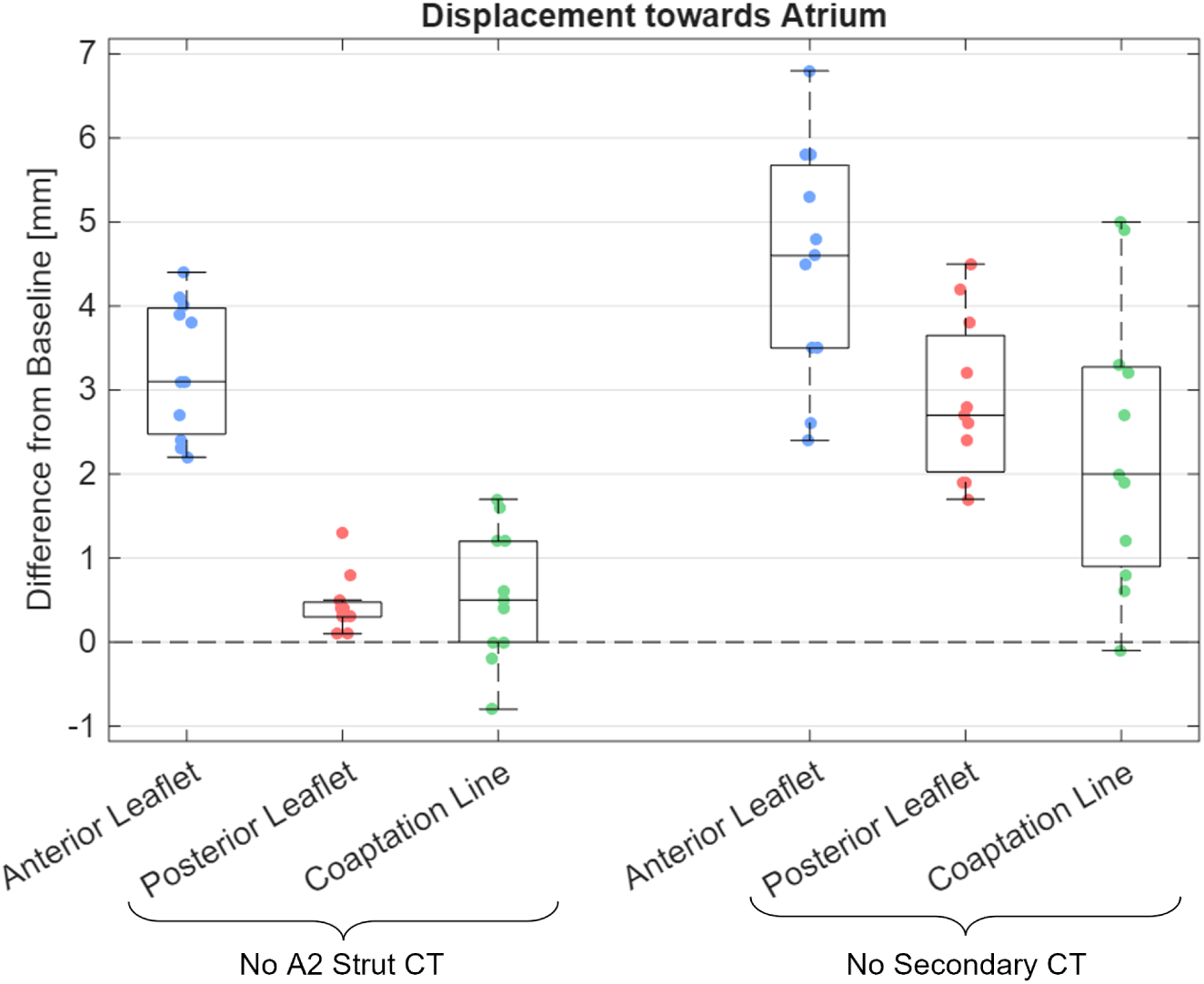
Billowing heights of individual leaflets and coaptation line displacement (equivalent to change in tenting height) toward the left atrium after transecting A2 strut chordae tendineae (CT) and all secondary CTs of the anterior and posterior leaflets.

### A.2 Statistics

For each comparison, a Shapiro–Wilk test was performed to assess normality of the data. Some datasets were normally distributed, while others were not, typically due to the presence of a single outlier. Given the small sample size (n = 11) and the occurrence of isolated outliers, it was considered justifiable to use either a paired t-test for normally distributed data or a Wilcoxon signed-rank test. The t-test is more sensitive when the normality assumption is met, whereas the Wilcoxon signed-rank test is more robust to deviations from normality.

To ensure that the choice of statistical test did not influence the interpretation of significance, all comparisons were evaluated using both methods. Ultimately, we chose to report the results of the Wilcoxon signed-rank test, as it is a median-based, nonparametric approach and therefore more appropriate for small sample sizes and data that may be affected by outliers.

An overview of the results of the Shapiro-Wilk tests, t-tests, Wilcoxon signed-rank tests, and the Bonferroni adjusted p-values is given in Table 1

**Table 1.** Statistical analysis of biomechanical parameters with Bonferroni adjusted p-values (3 tests per metric) of Wilcoxon signed-rank test. SW-Test: Shapiro-Wilk normality test (n = normal, nn = non-normal); T-test: paired t-test (OT = one tailed, TT = two tailed); Wilcoxon: Wilcoxon signed-rank test. Significance: ∗*p <* 0.05, ∗ ∗ *p <* 0.01, ∗ ∗ ∗*p <* 0.001.

| Comparison | Billowing Height |  |  |  |
| --- | --- | --- | --- | --- |
| | SW-Test | T-test (OT) | Wilcoxon | $p_{adj}$ |
| Baseline / A2 Strut CT | $n$ | $< 0.001^{***}$ | $< 0.001^{***}$ | $0.003^{**}$ |
| Baseline / All sec. CT | $n$ | $< 0.001^{***}$ | $< 0.001^{***}$ | $0.003^{**}$ |
| A2 Strut / All sec. CT | $n$ | $< 0.001^{***}$ | $< 0.001^{***}$ | $0.003^{**}$ |

| Comparison | Coaptation Height |  |  |  |
| --- | --- | --- | --- | --- |
| | SW-Test | T-test (TT) | Wilcoxon | $p_{adj}$ |
| Baseline / A2 Strut CT | $nn$ | $0.036^*$ | $< 0.001^{***}$ | $0.003^{**}$ |
| Baseline / All sec. CT | $n$ | $0.513$ | $0.567$ | $1.000$ |
| A2 Strut / All sec. CT | $n$ | $0.051$ | $0.042^*$ | $0.126$ |

| Comparison | Coaptation Area |  |  |  |
| --- | --- | --- | --- | --- |
| | SW-Test | T-test (TT) | Wilcoxon | $p_{adj}$ |
| Baseline / A2 Strut CT | $n$ | $0.0597$ | $0.117$ | $0.352$ |
| Baseline / All sec. CT | $nn$ | $0.0371^*$ | $0.010^{**}$ | $0.029^*$ |
| A2 Strut / All sec. CT | $nn$ | $0.046^*$ | $0.010^{**}$ | $0.029^*$ |

### A.3 Biomechanical parameters

Figure 10 gives an overview of the geometric features addressed in this manuscript and defined in the methods section.

**Fig. 10.**
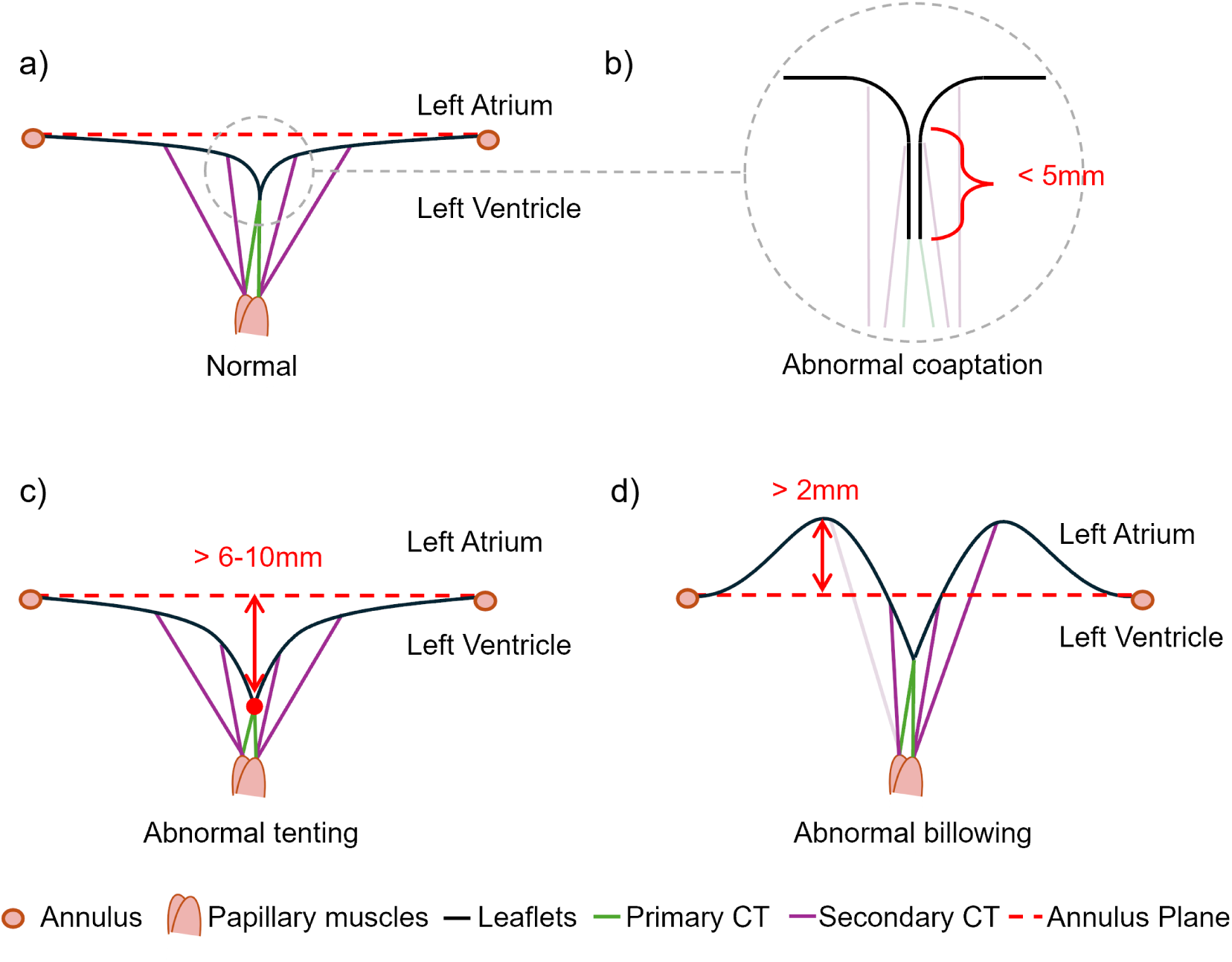
Schematic parasternal long-axis views of a) a normal mitral valve, b) the coaptation zone of an abnormal valve, defined by a coaptation height below 5 mm, c) an abnormal valve with tenting defined by a tenting height above 6 to 10 mm, and d) an abnormal valve with billowing defined by a billowing height greater than 2 mm. CT = chordae tendineae.

Table 2 presents all measured biomechanical parameters.

**Table 2.** Biomechanical parameters of porcine mitral valves under different conditions (n=11). CT = Chordae Tendineae, PLAX = Parasternal long-axis view. – Coaptation height decreased at PLAX already; * Coaptation height did not decrease at any of the ligating clip positions.

|  | MV1 | MV2 | MV3 | MV4 | MV5 | MV6 | MV7 | MV8 | MV9 | MV10 | MV11 | Median |
| --- | --- | --- | --- | --- | --- | --- | --- | --- | --- | --- | --- | --- |
| **Billowing Height (mm)** |  |  |  |  |  |  |  |  |  |  |  |  |
| Baseline / A2 Strut CT | 3.1 | 3.8 | 2.4 | 3.9 | 2.8 | 3.1 | 4.4 | 4.0 | 4.5 | 2.2 | 2.3 | **3.1** |
| Baseline / All sec. CT | 4.5 | 4.8 | 3.8 | 5.3 | 3.4 | 5.6 | 5.8 | 5.8 | 6.8 | 2.7 | 4.5 | **4.8** |
| A2 Strut CT / All sec. CT | 1.4 | 1.0 | 1.4 | 1.4 | 0.6 | 2.5 | 1.4 | 1.8 | 2.3 | 0.5 | 2.2 | **1.4** |
| **Coaptation Height at PLAX view (mm)** |  |  |  |  |  |  |  |  |  |  |  |  |
| Baseline / A2 Strut CT | -1.2 | -0.4 | -5.2 | -1.7 | -0.1 | -0.5 | -0.7 | -0.1 | -0.6 | -0.1 | -1.2 | **-0.6** |
| Baseline / All sec. CT | 3.3 | 0.6 | -5.1 | -0.8 | 0.0 | 4.5 | 4.2 | -2.7 | 1.4 | -0.9 | 2.1 | **0.6** |
| A2 Strut CT / All sec. CT | 4.5 | 1.0 | 0.1 | 0.9 | 0.1 | 5.0 | 4.9 | -2.6 | 2.0 | -0.8 | 3.3 | **0.9** |
| **Coaptation Height in other locations (mm)** |  |  |  |  |  |  |  |  |  |  |  |  |
| Baseline / All secondary CT | -0.9 | * | – | – | -0.6 | -5.8 | -1.8 | – | -2.6 | – | * |  |
| **Coaptation Area (cm<sup>2</sup>)** |  |  |  |  |  |  |  |  |  |  |  |  |
| Baseline / A2 Strut CT | -0.24 | 0.06 | -0.02 | -0.14 | 0.03 | 0.06 | 0.00 | -0.34 | -0.27 | -0.03 | -0.11 | **-0.03** |
| Baseline / All sec. CT | -3.19 | -2.95 | -3.26 | -3.82 | -3.18 | -3.12 | -3.33 | -6.04 | -3.66 | -3.72 | -3.35 | **-3.33** |
| A2 Strut CT / All sec. CT | -2.95 | -3.01 | -3.24 | -3.68 | -3.21 | -3.18 | -3.34 | -5.70 | -3.39 | -3.69 | -3.24 | **-3.24** |
| **Physiological Baseline (mm)** |  |  |  |  |  |  |  |  |  |  |  |  |
| Tenting | 0.4 | 2.2 | 1.5 | 0.8 | 4.9 | 1.4 | 1.1 | -1.2 | 1.2 | 1.2 | 3.9 | **1.2** |
| Billow height | 0.6 | 2.5 | 1.8 | 0.8 | 0.2 | 0.7 | 0.3 | 4.5 | 1.4 | 1.8 | 0.1 | **0.8** |
| **Individual Leaflet Billowing Height vs Coaptation Line Displacement (=change in tenting height) (mm)** |  |  |  |  |  |  |  |  |  |  |  |  |
| Anterior leaflet (A2 Strut CT) | 3.1 | 3.8 | 2.4 | 3.9 | 2.7 | 3.1 | 4.4 | 4.0 | 4.1 | 2.2 | 2.3 | **3.1** |
| Posterior leaflet (A2 Strut CT) | 0.1 | 0.3 | 0.1 | 0.3 | 0.3 | 0.8 | 0.4 | 1.3 | 0.5 | 0.4 | 0.3 | **0.3** |
| Coaptation line (A2 Strut CT) | -0.8 | 0.5 | -0.2 | 1.2 | 0.4 | 1.7 | 1.6 | 0.6 | 0.0 | 1.2 | 0.0 | **0.5** |
| Anterior leaflet (All CT) | 4.5 | 4.8 | 2.6 | 5.3 | 3.5 | 4.6 | 5.8 | 5.8 | 6.8 | 2.4 | 3.5 | **4.6** |
| Posterior leaflet (All Strut CT) | 3.2 | 1.7 | 3.8 | 1.9 | 1.9 | 2.8 | 4.2 | 2.4 | 2.6 | 2.7 | 4.5 | **2.7** |
| Coaptation line (All Strut CT) | 1.9 | 2.0 | -0.1 | 3.3 | 0.8 | 4.9 | 5.0 | 0.6 | 3.2 | 1.2 | 2.7 | **2.0** |

